# Acute Local and Systemic Effects of Intra-Articular Administration of Liposomal IDO-1 following Anterior Cruciate Ligament Rupture

**DOI:** 10.64898/2025.12.09.693103

**Authors:** Mackenzie Fleischer, Michael Newton, Samantha Hartner, Christopher J. Bush, Anthony Arveschoug, Christopher Vasileff, Erin A. Baker, Kevin C. Baker

## Abstract

Joint injuries, such as rupture of the anterior cruciate ligament (ACL), is associated with the development of post-traumatic osteoarthritis (PTOA). It is known that ACL rupture can lead to disruption of metabolic pathways, including the conversion of the essential amino acid tryptophan to kynurenine, which is associated with a sustained inflammatory response. An in vivo study was undertaken to determine the acute effects of intra-articular administration of liposomes loaded with the tryptophan-catabolizing enzyme indoleamine 2,3-dioxygenase-1 (IDO-1) following ACL rupture. Using an established rat model of non-surgical ACL injury, male and female Lewis rats underwent a single intra-articular injection of empty liposomes, or liposomes loaded with IDO-1 and were subsequently randomized to 1- or 2-week endpoints. IDO-1 treatment after ACL injury was associated with a significant reduction in synovial fluid concentration of tryptophan at both 1-and 2-week endpoints. In addition to a reduction in tryptophan, IDO-1 treatment led to significantly lower synovial fluid concentrations of IL-1b and TNF-a. Intra-articular administration of IDO-1-loaded liposomes also increased the ratio of regulatory T lymphocytes (Tregs) to IL-17-secreting helper T lymphocytes (Th17 cells). Similarly, IDO-1 treatment increased the number of CTLA4+ cells relative to IL-17A+ cells that infiltrated joint tissues at a 2-week endpoint. Contrast-enhanced micro-computed tomography (CE-uCT) was used to quantify treatment-based effects on articular cartilage thickness and surface roughness at at 2-week endpoint. In addition to sex-based differences, IDO-1-loaded liposome treatment was associated with increased cartilage thickness, with no significant effects on surface roughness. Histologic characterization is needed to determine whether this increased cartilage thickness represents a chondroprotective effect, or a degenerative effect of IDO-1-treatment.

## INTRODUCTION

The anterior cruciate ligament (ACL) is a primary stabilizing structure of the knee joint, which resists translation of the tibia. Rupture of the ACL initiates a complex inflammatory cascade resulting in significant pain and debility. This inflammatory cascade has also been implicated in the development of post-traumatic osteoarthritis (PTOA).^1–6^ PTOA is characterized by progressive degeneration of bone and articular cartilage, which is attributed to a specific episode of joint trauma.^7,8^ Studies have observed that, without surgical intervention, up to 90% of people experiencing an ACL rupture will develop PTOA.^5^ Despite advances in surgical technique and technology, more than half of patients that opt for ACL reconstructive surgery will ultimately develop PTOA, suggesting that acute biologic processes may be responsible for disease onset and progression.^1,3^ This concept has been corroborated in animal models of “idealized” ACL reconstruction, where detachment of the ACL from its insertion followed by immediate reattachment, was associated with the development of osteoarthritic changes.^9–12^ Due to high rates of PTOA after ACL injuries, there is significant interest in the development of therapeutic strategies to interfere with these acute processes to mitigate ACL rupture-induced PTOA.

Following ACL injury, tissue and synovial fluid increases in pro-inflammatory cytokines, matrix-degrading enzymes, and other soluble mediators have been observed.^1–3,13,14^ Upstream, there are alterations in cell physiology, including the disturbance of metabolic pathways.^7,11^ Metabolomic profiling has identified tryptophan metabolism as a dysregulated metabolic pathway in numerous inflammatory arthropathies, musculoskeletal disorders, and ACL rupture-induced PTOA.^15–20^ More than 95% of tryptophan is catabolized via the Kynurenine Pathway by enzymes indoleamine 2,3-dioxygenase-1 (IDO-1), indoleamine 2,3-dioxygenase-2 (IDO-1), or tryptophan 2,3-dioxygenase (TDO).^21^ Conversion of tryptophan to kynurenine by these enzymes results in transient activation of aryl hydrocarbon receptor (AhR) signaling, which is associated with expansion of immunomodulatory cell populations, including regulatory T lymphocytes (T_regs_).^22–27^ Decreased conversion of tryptophan to kynurenine, which has been observed in animal models of ACL rupture-induced PTOA, is associated with increased inflammation and expansion of inflammatory cell types, including IL-17-secreting helper T lymphocytes (T_h_17).

The stalled conversion of tryptophan to kynurenine, which has been observed following ACL rupture, represents a potential therapeutic target for the mitigation of PTOA. We designed and performed this study to assess the acute effects of the intra-articular administration of IDO-1 on tryptophan metabolism, inflammation, circulating and infiltrating T lymphocyte subsets, as well as articular cartilage structure. The primary objective of this study was to determine if local administration of IDO-1 would alter the local inflammatory response to ACL rupture, by promoting the conversion of tryptophan to kynurenine and downstream kynurenine metabolites.

## METHODS

### Liposomal Encapsulation of IDO-1

To quantitatively evaluate IDO-1 encapsulation efficiency and release profile, the stockprotein was fluorescently tagged prior to experimentation. Recombinant murine IDO-1 protein (R&D Systems Inc.) was labeled using Alexa Fluor™ 488 Microscale Protein Labeling Kit (ThermoFisher), according to manufactures instructions. Briefly, 20µg of IDO-1 was reconstituted in DPBS at a concentration of 1mg/mL. The stock IDO-1 was pH-adjusted using sodium bicarbonate prior to adding 1.2μL of reactive dye. The reaction was then incubated for 15 minutes at room temperature. Unreacted dye was removed from the solution via spin column filtration. The purified dye-labeled protein was collected and spectrophotometrically analyzed to determine the resulting protein concentration and degree of labeling.

Commercially-available, empty dipalmitoyl phosphatidylcholine (DPPC) liposomes (NOF America) were used as a vehicle for intra-articular IDO-1 delivery. Liposomes were stored as a lyophilized cake prior to experimentation. Liposome formation and protein loading occurs spontaneously as a dehydration-rehydration reaction upon addition of an aqueous solution of the target drug to the lyophilized DPPC cake. Based on IDO-1 structure and molecular charge, the cationic formulation was used to increase encapsulation. Liposomes were formed by adding 2.0mL of deionized sterile water containing the desired dose of IDO-1 followed by gentle inversion of the mixture. DPPC liposomes were loaded with 10, 100, or 1000ng of IDO-1 within a 50μL total volume per dose.

Liposomes were formed at each concentration of fluorescently-tagged IDO-1, using the described protocol. The liposomes were then loaded into a disposable dialysis device with a 50kD molecular weight cut-off to remove unencapsulated protein while retaining intact liposomes. After 2 hours, the liposomes were retrieved and transferred into a 96-well plate with 50μL per well. IDO-1 concentration was calculated via fluorescent intensity detection. Known concentrations of fluorescently-tagged IDO-1 were used to generate a standard curve and empty (not loaded) liposomes were used as a control. The initial dose was then adjusted accordingly to achieve the desired dose and a full-scale experiment was conducted to ensure repeatability.

### In Vitro Tryptophan Conversion Assay

To assess whether DPPC liposome encapsulation of IDO-1 interferes with tryptophan-metabolizing functionality, an *in vitro* IDO-1 activity assay was performed. The assay measured IDO activity via L-tryptophan metabolism to N-formylkynurenine (NFK). The assay was conducted according to manufacturer’s instructions (Sigma Aldrich). Briefly, IDO-1-loaded liposomes at each dose (10, 100, 1000ng) were incubated with L-tryptophan. After 45 minutes, a fluorogenic developer that selectively reacts with NFK was added to each sample. After 3 hours, the fluorescence was read at 488nm. NFK concentrations in each sample were calculated from a standard curve and used to derive IDO-1 metabolic activity. Samples containing empty liposomes or an IDO-1 inhibitor were used as controls.

### ACL Rupture Procedure and Intra-Articular Administration of IDO-1-Loaded Liposomes

In accordance with an IACUC-approved protocol, 14-week-old male and female Lewis rats were anesthetized and underwent a non-surgical ACL rupture, as previously described.^28–30^ Briefly, rats were anesthetized via inhaled isoflurane and placed prone on an environmentally-controlled stage. The right knee was flexed to 100° and the foot was placed into a jig attached to the actuator of an electromechanically-actuated materials testing frame (Insight 5, MTS Systems Inc.) After applying a 3N preload and sinusoidal preconditioning loads (1-5N), a rapid 3.0mm displacement was applied to the right tibia via the actuator resulting a closed, isolated ACL rupture. Animals received a subcutaneous injection of long-acting buprenorphine (BuprenexSR, 1.2mg/kg) for post-procedural analgesia. Sham animals underwent identical anesthesia and analgesia protocols. They were placed on the materials testing frame and underwent both the preload and preconditioning cycles, but were not subjected to the rapid tibial displacement. All rats were allowed *ad libitum* activity with 12-hour light/dark cycles and free access to food and water.

Twenty-four hours after ACL rupture (or Sham), animals were randomized to receive intra-articular injections of IDO-1-loaded DPPC liposomes (10, 100, or 1000ng total dose) or Control (empty DPPC liposomes.) Rats were anesthetized with via inhaled isoflurane, and then the right limb was shaved and sterile-prepped with alternating scrubs of iodine and chlorhexidine gluconate solutions. Using a 28G needle attached to a microsyringe, 50mL of liposome solution (IDO-1-loaded or empty) was injected into the right knee and the joint was gently ranged several times after withdrawing the needle. Animals were then recovered from anesthesia and allowed to return to *ad libitum* activity.

### Flow Cytometry of Whole Blood

To assess systemic effects of intra-articular IDO-1-loaded liposome treatment, flow cytometry was performed. Whole blood collected at 1- and 2-week endpoints underwent red blood cell lysis followed by magnetic bead-assisted cell isolation of T lymphocytes (Pan T Cell Isolation Kit – Rat, Miltenyi Biotec). The isolated T cell population was then labeled for flow cytometric evaluation to further characterize systemic T cell phenotype shifts between IL-17-secreting helper T cell (T_h_17) and FOXP3^+^ regulatory T cell (T_reg_) populations. Cells were labeled with antibodies against CD4 to confirm T-cell linage, IL-17 to identify Th-17 cells, and CD25 with FOXP3 to identify T_reg_ cells. Samples were run in duplicate on a multi-color flow cytometer. The T_reg_/T_h_17 ratio was calculated by dividing the percentage of CD4^+^, CD25^+^, and FOXP3^+^ cells by percentage of CD4^+^ and IL17^+^ cells. The ratio was compared between groups was compared at both the 1-and 2-week endpoints, using a one-way ANOVA with post-hoc Dunnett’s tests (p<0.05) for multiple comparisons.

### Quantification of Synovial Fluid Tryptophan and Inflammatory Cytokines

Synovial fluid was aseptically aspirated from both ACLR and contralateral joints for analysis of tryptophan, kynurenine, IL-1β, and TNF-α concentration via ELISA. Analyte concentrations were normalized to protein content, as quantified by a commercially-available bicinchonic acid assay. Synovial fluid concentrations of tryptophan and kynurenine were quantified at both the 1- and 2-week endpoint and compared between treatment groups (0, 10, 100, or 1000ng of IDO-1-loaded liposomes), using a one-way ANOVA with post-hoc Tukey’s tests (p<0.05) for multiple comparisons. Synovial fluid concentrations of IL-1β and TNF-α were quantified at the 1-week endpoint and compared between treatment groups, using a one-way ANOVA with post-hoc Tukey’s test (p<0.05) for multiple comparisons.

### Contrast-Enhanced Micro-Computed Tomography of Knee Joints

Two weeks after ACL rupture and intra-articular treatment, knee joints were aseptically harvested *postmortem*. Whole joints were incubated in a 20% v/v solution of ioxaglate (Hexabrix 320, Guerbet LLC, Bloomington, IN) in PBS containing protease inhibitors at room temperature for 2 hours, as previously described.^32^ Samples were imaged via μCT (MicroCT-40, Scanco Medical, Zurich, Switzerland) at 55kvP, 145μA, and a 320ms integration time with an 8μm isotropic voxel size. Hydroxyapatite phantoms of known mineral density were used for calibration. Exported DICOM stacks underwent semi-automated segmentation of distal femoral articular cartilage (AC), guided by an experienced researcher using a validated, atlas-based approach.^33^ Parameterized 2D mappings of the femoral AC surface were generated, and then used to quantify articular cartilage thickness (AC.Th) and areal surface roughness (AC.Sa) on the medial femoral condyle, lateral femoral condyle, patellofemoral compartment (trochlea), and globally for the entire distal femoral articular surface, as previously described.^31^ AC.Th and AC.Sa measurements in each compartment were compared between groups (empty liposomes vs. 100ng IDO-1-loaded liposomes), using a one-way ANOVA with post-hoc Tukey’s tests (p<0.05) for multiple comparisons.

### Histology and Immunohistochemistry of Knee Joints

After serial rinses in PBS to remove the contrast agent, limbs were fixed in 10% neutral buffered formalin. Joints were decalcified in EDTA and embedded in paraffin for sectioning. Blocks were sectioned coronally (4μm sections) and stained with Hematoxylin & Eosin and immunostained for IL-17A (T_h_17 marker) and CTLA4 (T_reg_ marker). Whole slides were digitally scanned at a magnification of 40x and analyzed by blinded reviewers.

## RESULTS

### Encapsulation and Release Kinetics of IDO-1 from Liposomes

IDO-1 was efficiently incorporated into DPPC liposomes at doses ranging from 10ng to 1000ng. Microscopy of loaded liposomes demonstrated that fluorescently-tagged IDO-1 protein was primarily incorporated within the lipid bilayer of the liposomes (**Figure 1A**). At the 10ng target dose (0.0002μg/μL), 57.8% of IDO-1 was successfully incorporated into the liposomes. Encapsulation efficiency peaked at the 100ng target dose (0.002μg/μL), with 97% of IDO-1 incorporated, and then decreased to 76% of IDO-1 incorporated at the 1000ng target dose (0.02μg/μL) (**Figure 1B**). These results were subsequently used to produced liposomes solutions containing specific doses (10, 100, or 1000ng) of IDO-1 into a 50µL solution for other *in vitro* and *in vivo* studies.

**Figure 1.**
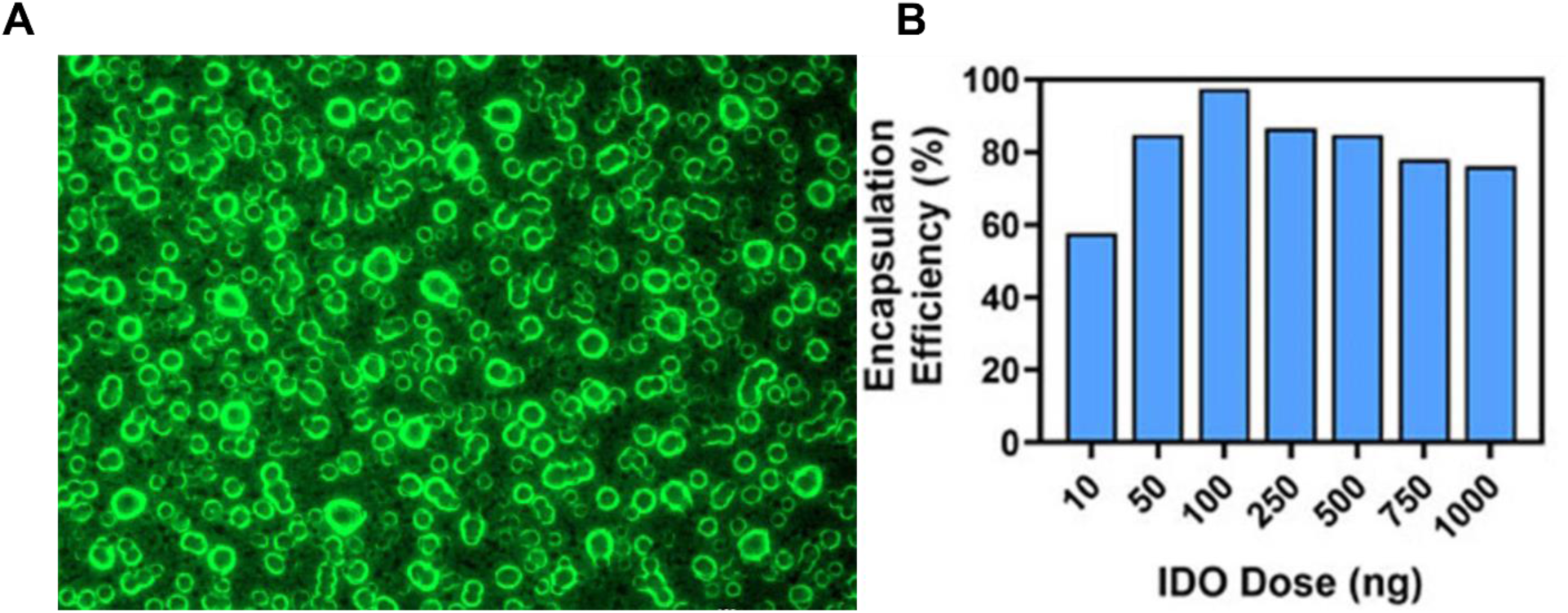
Fluorescence microscopy demonstrating successful incorporation of AlexaFluor 488-tagged IDO-1 into DPPC liposomes (A). Encapsulation efficiency peaked at a target dose of 100ng, with 97% of IDO-1 incorporated into liposomes (B).

### Tryptophan Catabolism by IDO-1-loaded Liposomes

*In vitro* assays were used to determine the impact of liposomal encapsulation on IDO-1 enzymatic activity. IDO-1 activity was assessed by measuring the conversion of L-tryptophan (Trp) to the metastable Kynurenine intermediate N-formylkynurenine (NFK). In addition to liposomal IDO, free IDO of the same concentrations was also assayed for comparison. Overall, the metabolic activity of IDO increased with increased dose concentration, with 10ng of liposomal IDO-1 yielding 168pmol NFK, and 10ng free IDO yielding 124pmol NFK. The 100ng liposomal IDO dose yielded 1,320pmol NFK, compared to the 100ng free IDO dose, which yielded 2,660pmol NFK. Lastly, the 1000ng liposomal IDO dose yielded 9,679pmol NFK, while the 1000ng free IDO dose yielded 32,955pmol NFK. The specific activity of IDO at each dose, defined by “one unit of IDO-1 activity is the amount of enzyme that generates 1μmol of detected N-formylkynurenine per minute by oxidative metabolism of 1μmol L-tryptophan at 37 °C,” was then calculated. The specific activities of the 10ng liposomal and free IDO doses were 374μU/mg and 275μU/mg, respectively. For the 100ng liposomal and free IDO doses, specific activities were 293μU/mg and 591μU/mg, respectively. Finally, the specific activity of the 1000ng liposomal IDO dose was 215μU/mg, while the specific activity of the 1000ng free IDO dose was 732μU/mg.

### Liposomal IDO-1 Reduces Intra-Articular Tryptophan Concentration

Following the confirmation of *in vitro* Tryptophan-catabolizing activity of IDO-1-loaded DPPC liposomes, a subsequent study of the *in vivo* Tryptophan-reducing capacity of the liposomes was assayed. In the Control (empty) group, there was an increase in Tryptophan concentration in synovial fluid from 1 week to 2 weeks post-ACL rupture. At 1 week post-injury, animals that received 1000ng of liposomal IDO-1 24 hours after ACL injury had significantly lower synovial fluid concentrations of Tryptophan (**Figure 2**). At 2 weeks post-injury, animals treated with 100ng or 1000ng of liposomal IDO-1 showed significantly lower synovial fluid concentrations of Tryptophan, compared to animals that received empty liposomes after ACL rupture.

**Figure 2.**
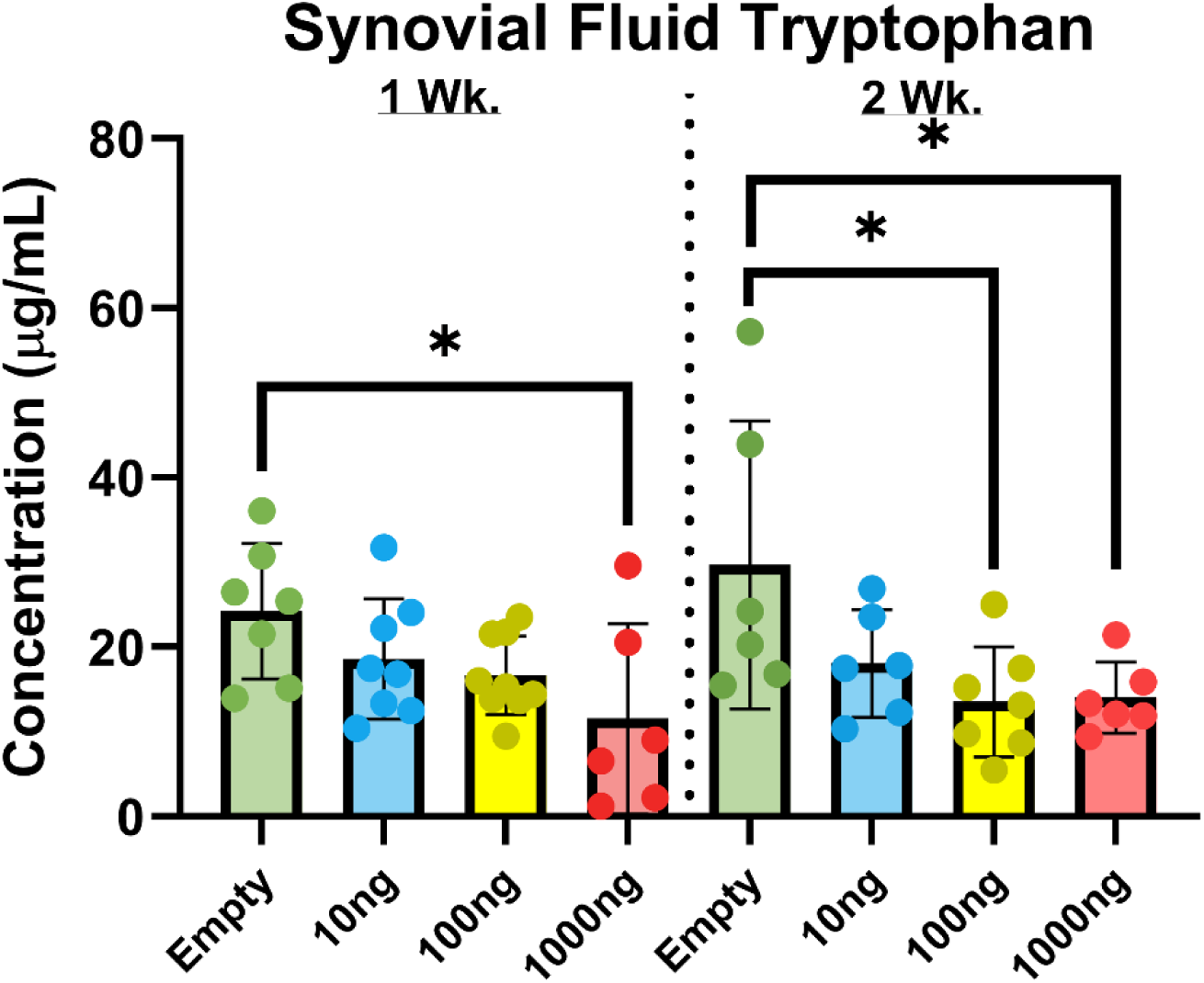
Significant reductions in synovial fluid concentrations of Tryptophan were observed following intra-articular administration of IDO-1-loaded liposomes after ACL rupture.

### Liposomal IDO-1 Reduces Intra-Articular Inflammatory Cytokines

In addition to intra-articular Tryptophan metabolism, the effect of liposomal IDO-1 on synovial fluid concentrations of inflammatory cytokines was investigated. With a single intra-articular injection of liposomal IDO-1 given 24 hours after ACL injury, both 10ng and 100ng doses led to significantly decreased concentrations of TNF-α (**Figure 3A**) at 2 weeks post-injury. The 10ng IDO-1 dose also led to decreased synovial fluid concentrations of IL-1β (**Figure 3B**) at 2 weeks post-injury.

**Figure 3.**
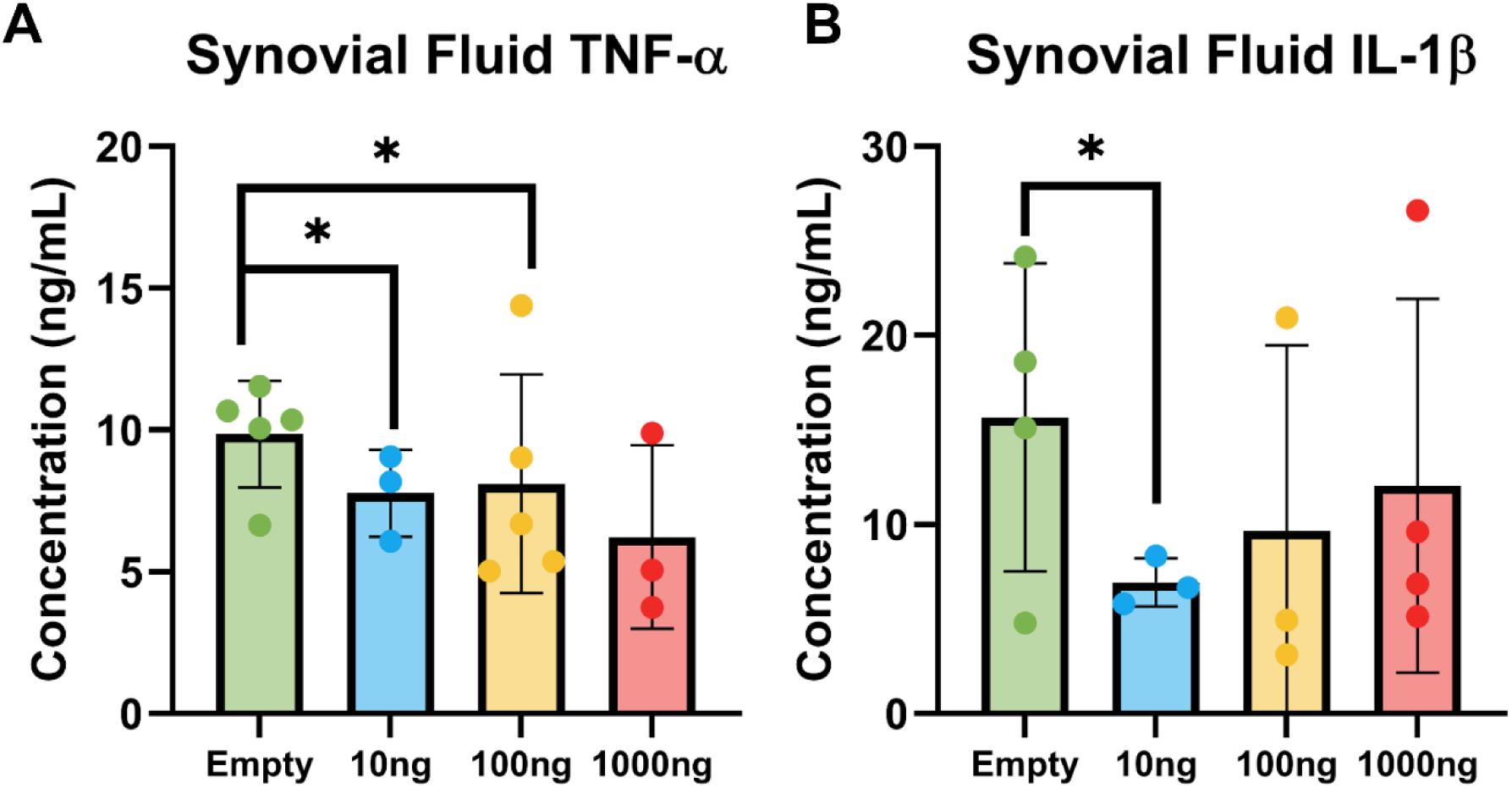
IDO-1-loaded liposomes administered intra-articularly 24 hours after ACL ruptures led to significantly reduced synovial fluid concentrations of TNF-α (A) and IL-1β (B), compared to animals that received empty liposomes.

### Effect of Intra-Articular IDO-1 Treatment on Circulating T Lymphocyte Profile

Circulating immune cell profiles have been correlated with disease state and progression in numerous inflammatory and musculoskeletal disorders, including OA.^34–35^ The balance of regulatory T lymphocytes (T_reg_) to IL-17-expressing helper T lymphocytes (T_h_17) is used to identify systemic shifts in anti-inflammatory versus pro-inflammatory activity.^36^ Intra-articular injection of IDO-1-loaded liposomes following ACL rupture was associated with an alteration of the T_reg_/T_h_17 ratio. Specifically, treatment with 100ng of IDO-1-loaded liposomes led to significantly increased T_reg_/T_h_17 ratio when compared to animals that received empty liposomes after ACL rupture (p=0.044, **Figure 4**).

**Figure 4.**
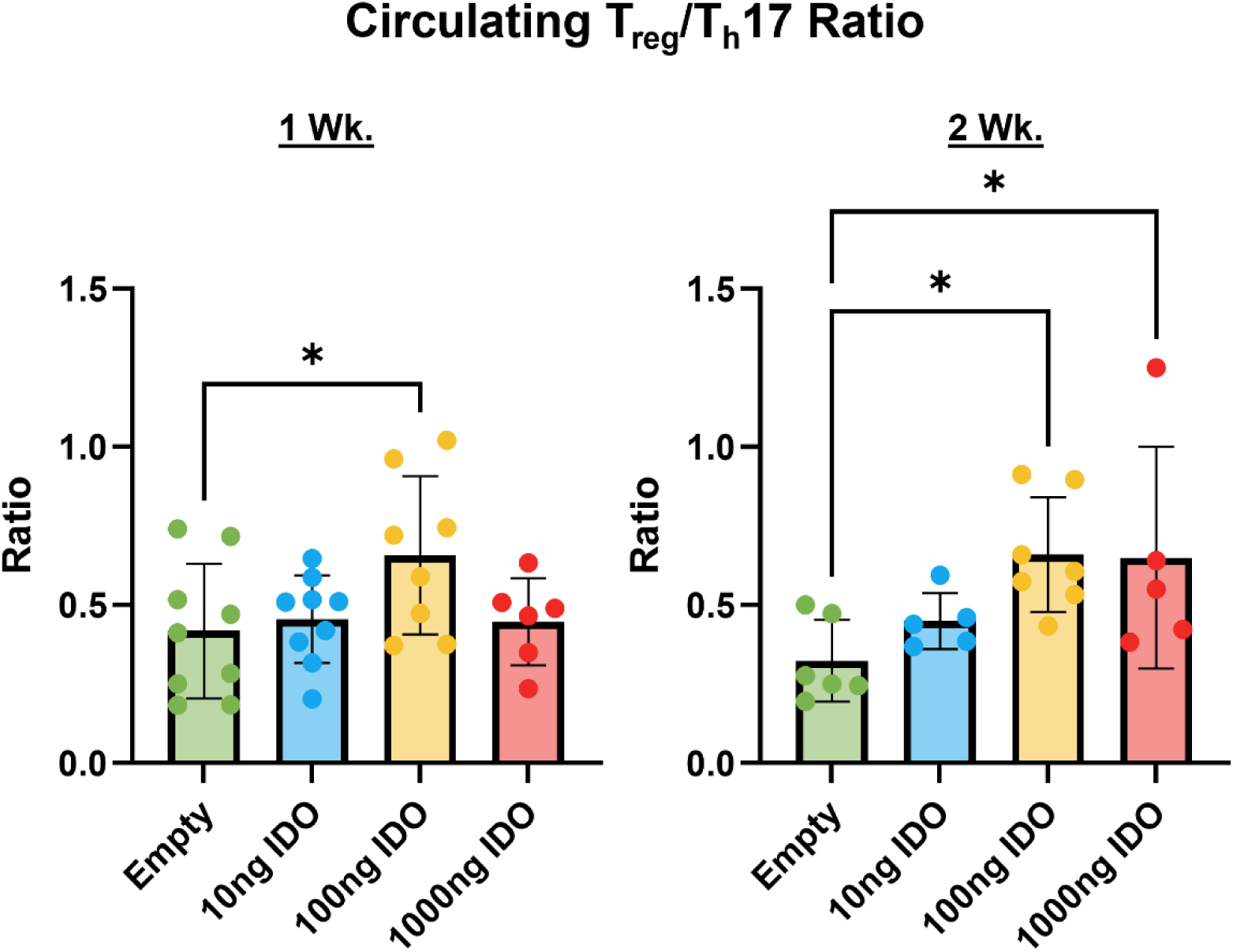
Comparison of the ratio of circulating T_regs_ (CD4^+^, CD25^+^, and FOXP3^+^) to T_h_17 cells (CD4^+^ and IL17^+^) between rats treated with intra-articular IDO-1-loaded liposomes following ACL rupture at 1 week (A) and 2 weeks (B) post-treatment. In Figure 4A, * denotes a statistically significant difference between the group treated with 100ng of IDO-1 vs. empty liposomes (p=0.044). In Figure 4B, * denotes statistically significant differences between the group treated with 100ng of IDO-1 vs. empty liposomes (p=0.022), and between the group treated with 1000ng of IDO-1 vs. empty liposomes (p=0.044).

### Effect of IDO-1 Treatment on Tissue Infiltration by T_regs_ and T_h_17 Lymphocytes

In addition to increasing the T_reg_/T_h_17 ratio of lymphocytes in circulation, treatment with IDO-1-loaded liposomes also influenced the composition of cells that infiltrated joint tissues after ACL rupture. Animals treated with IDO-1-loaded liposomes demonstrated more cells that stained positive for the immune checkpoint molecule CTLA-4, when compared to animals that received empty liposomes (**Figure 5**). These positive-staining cells were located predominatly within the intracondylar notch at the femoral insertion of the ACL, with intense staining noted around the genicular artery. Conversely, animals treated with IDO-1-loaded liposomes showed substantially less IL-17A staining compared to animals treated with empty liposomes (**Figure 6**). IL-17A staining in the empty liposome-treated animals was most intense in the synovium.

**Figure 5.**
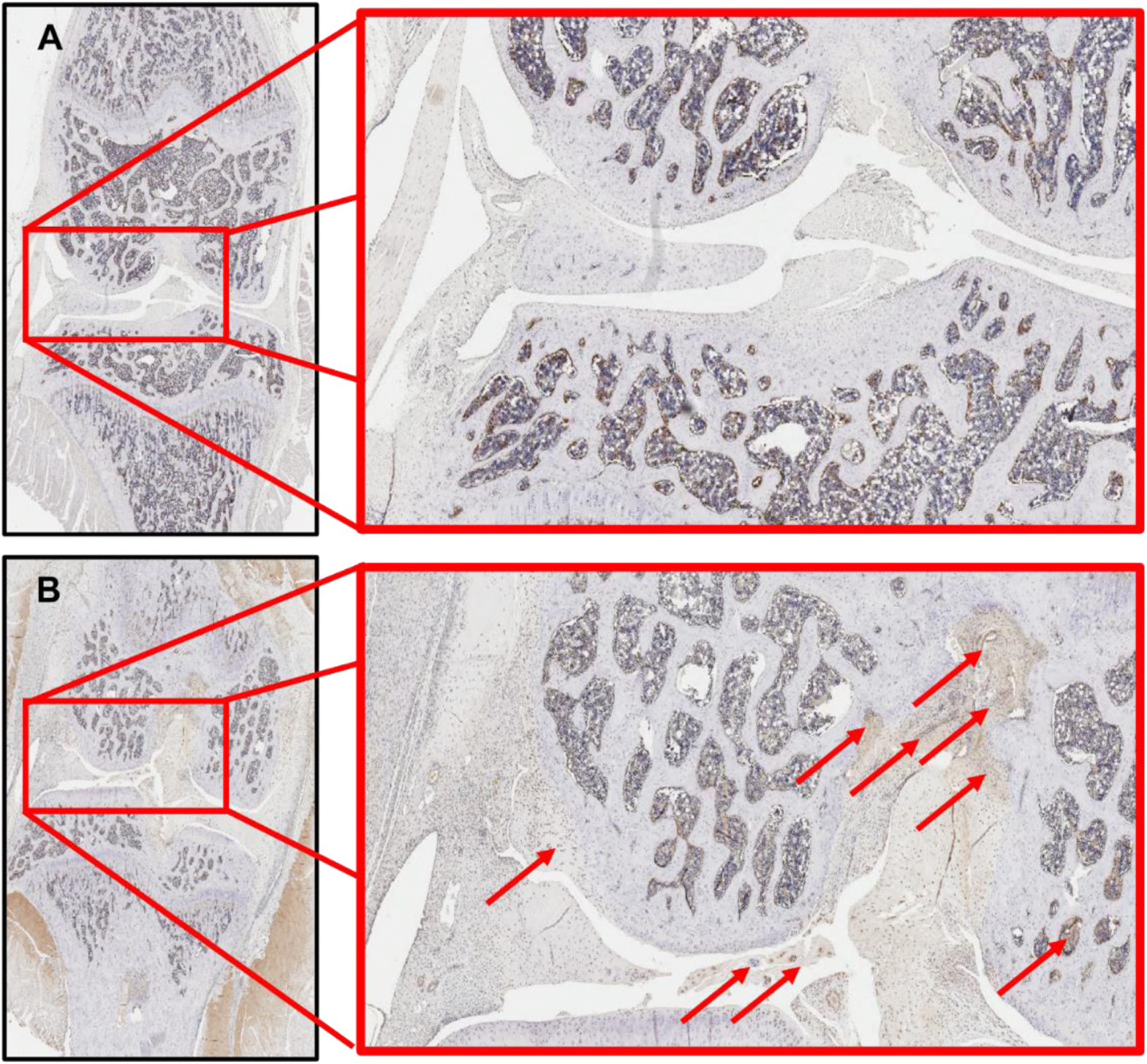
Coronal sections of ACL-injured and treated knees immunostained with CTLA-4. Joint tissues from animals treated with empty liposomes (A) display minimal staining, while joint tissue from animals treated with 100ng IDO-1 liposomes (B) show more positively-stained cells, particularly in the intracondylyar notch at the insertion of the ACL.

**Figure 6.**
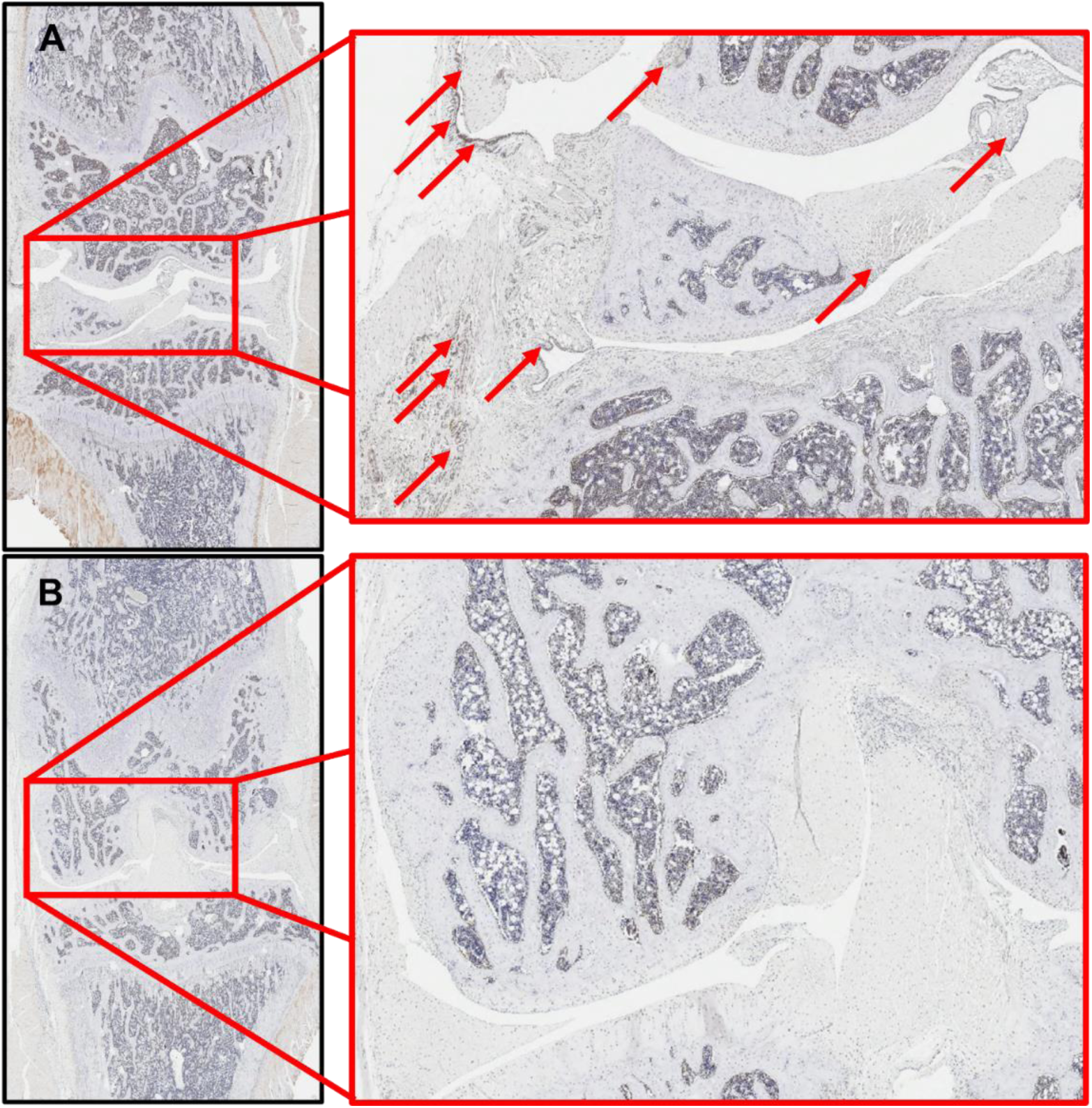
Coronal sections of ACL-injured and treated knees immunostained with IL-17A. Joint tissues from animals treated with empty liposomes (A) display staining in synovium and other peri-articular structures, while joint tissue from animals treated with 100ng IDO-1 liposomes (B) show substantially less staining throughout the joint.

### Acute Effects of IDO-1 Therapy on Articular Cartilage

CE-μCT imaging was used to quantify thickness (AC.Th) and areal surface roughness (AC.Sa) of the distal femoral articular cartilage of both right (ACL-injured) and left (uninjured) femora in animals that received intra-articular injections of empty liposomes (Empty) or liposomes filled with 100ng of IDO-1 (100ng IDO). Significant sex-based differences in AC.Th and AC.Sa were observed in most compartments (**Figure 7**). Female rats treated with IDO-1-loaded liposomes demonstrated significant increases in AC.Th on both the lateral femoral condyle and the trochlea, as well as globally across the whole distal femoral articular surface when compared to ACL-injured animals treated with empty liposomes. Whole femur AC.Th was also higher in female rates treated with IDO-1-loaded liposomes compared to their own contralateral (uninjured) limbs. Male rats treated with IDO-1-loaded liposomes showed a significant increase in AC.Th on the medial femoral condyle compared to their contralateral (uninjured) limb, but no difference compared to male rats with ACL-injured limbs treated with empty liposomes. Whole femur and trochlear AC.Sa were significantly higher in for male rats compared to female rats, however, no statistically significant effects of IDO-1 treatment were observed.

**Figure 7.**
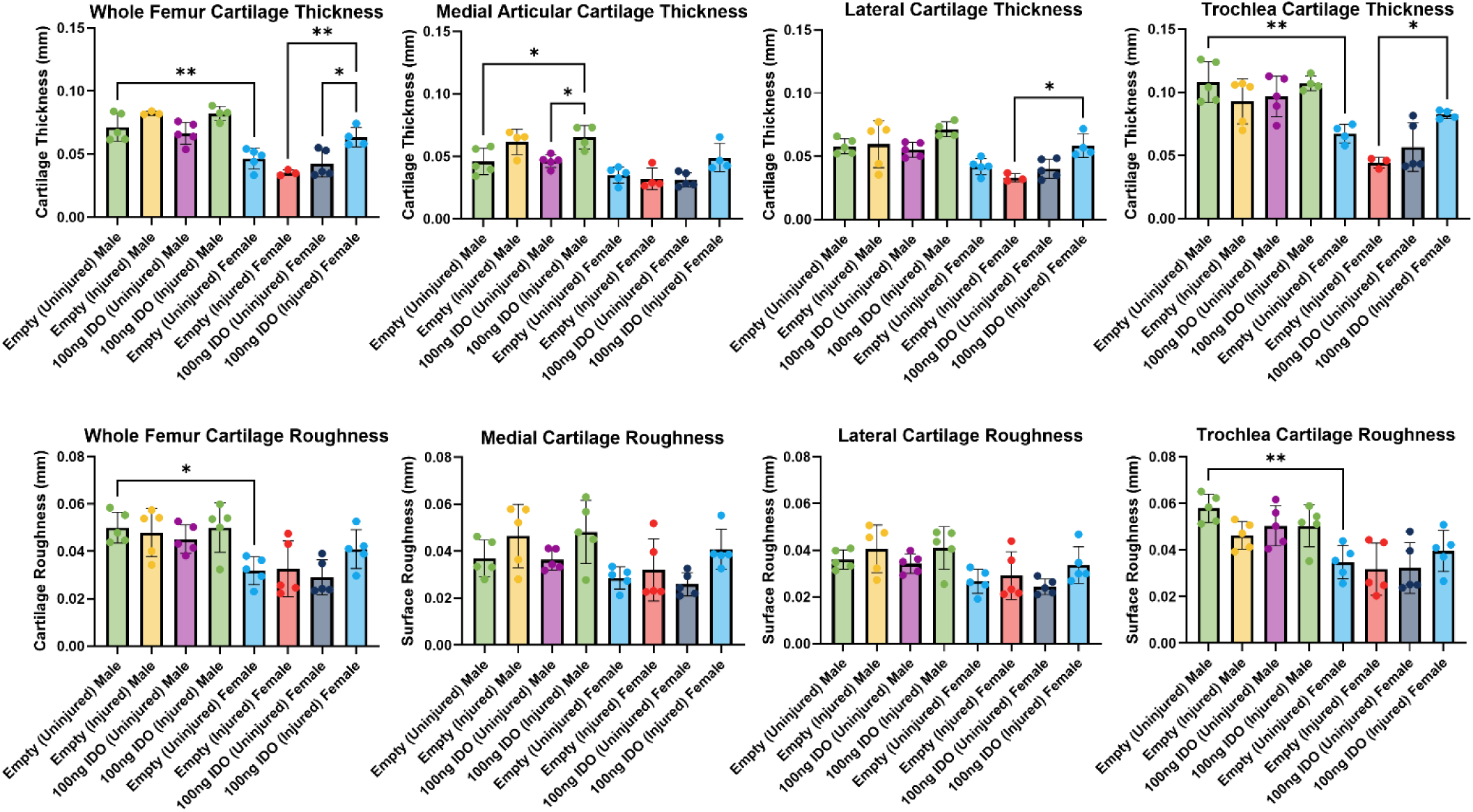
Articular cartilage thickness (AC.Th, Top Row) and areal surface roughness (AC.Sa) of articular cartilage (Bottom Row) on whole femur, medial femoral condyle, lateral femoral condyle, or trochlea, as quantified by CE-μCT. These data represented both the ACL-ruptured (Injured) and contralateral (Uninjured) limbs of animals that received either empty liposomes (Empty) or 100ng of IDO-1-loaded liposomes (IDO-1). One-way ANOVA with a post-hoc Tukey’s test for multiple comparisons, where * is p<0.05, and ** is p<0.001.

## DISCUSSION

Previous studies have reported that the conversion of the essential amino acid, Tryptophan, to Kynurenine as dysregulated in numerous inflammatory disorders.^15–18^ Similarly, traumatic joint injury, such as an ACL rupture, has been associated with a sustained decrease in Tryptophan catabolism.^20^ The focus of this study was to investigate whether increasing Tryptophan catabolism via intra-articular delivery of the enzyme, IDO-1, would affect the systemic and local inflammatory response to joint trauma.

In this study, IDO-1 was encapsulated in dipalmitoyl phosphatidylcholine (DPPC) liposomes via dehydration-rehydration reaction, with an efficiency exceeding 96% using a 100ng enzyme dose. Encapsulation of therapeutics for intra-articular delivery is a common practice due to the harsh microenvironment of synovial joints following injury.^37^ Liposomal systems have a multi-decade history as delivery agents for intra-articular therapeutics, such as corticosteroids and local anesthetics.^38,39^ DPPC-based liposomes have demonstrated biocompatibility within the knee joint in both animal models and humans.^40,41^ *In vitro* Tryptophan catabolism assays demonstrated that DPPC liposome-encapsulated IDO-1 converted Tryptophan substrate to the metastable intermedial, n-Formyl kynurenine. The specific activity of free IDO-1 was greater than that of the encapsulated IDO-1 suggesting that, in an *in vitro* setting, the liposomes may influence the rate of Tryptophan catabolism. Despite this potential interference, *in vivo* assays demonstrated a significant, dose-dependent reduction of Tryptophan concentration in synovial fluid at both 1 and 2 weeks post-ACL rupture, with intra-articular administration of liposomal IDO-1 (**Figure 2**).

In addition to reducing synovial fluid concentrations of Tryptophan, animals treated with liposome-encapsulated IDO-1 showed a significant decrease in synovial fluid concentration of the inflammatory cytokines, TNF-α and IL-1β, at 1 week post-ACL rupture. This observation aligns with previous studies that have shown that joint trauma, such as an ACL injury, has been associated with a significant increase in synovial fluid levels of pro-inflammatory cytokines, including TNF-α and IL-1β.^1–3,7–9^ Dysregulated tryptophan catabolism may also promote inflammation through impaired activation of the aryl hydrocarbon receptor (AhR).^42^ Intra-articular administration of liposomal IDO-1 promotes the catabolism of Tryptophan to both Kynurenine and downstream kynurenine metabolites, which suppress inflammation via activation of AhR signaling.^43–45^

Tryptophan metabolism also influences inflammation by altering the suppression or expansion of T lymphocytes. Specifically, the catabolism of Tryptophan to Kynurenine activates AhR signaling, which suppresses pathogenic T_h_17 differentiation and expands T_reg_ pools.^22–27^ In this study, intra-articular administration of a single-dose of IDO-1-loaded liposomes following ACL rupture was associated with a significantly higher ratio of T_regs_ (CD4^+^, CD25^+^, and FOXP3^+^) to T_h_17 lymphocytes (CD4^+^ and IL-17^+^), as quantified by flow cytometry of whole blood. This effect was sustained, as the T_reg_/T_h_17 ratio was greater in animals treated with either 100ng or 1000ng IDO-1 doses, compared to empty liposomes, at 2 weeks post-ACL injury and treatment. The ratio of circulating T_regs_ and T_h_17 cells has been correlated with disease activity in various immune-mediated conditions, with an increase in T_h_17 cells relative to T_regs_ associated with increased severity of symptoms.^46–48^ In addition to this shift in systemic T lymphocyte profile, immunostaining showed qualitative differences in the number of CTLA4^+^ and IL-17A^+^ cells infiltrating knee joint tissues. CTLA4 is an immune checkpoint molecule commonly expressed by T_regs_, while IL-17A is commonly expressed by Th17 cells. Animals treated with IDO-1-loaded liposomes demonstrated a greater number of CTLA4+ cells, when compared to animals that received empty liposomes. These CTLA4+ cells were located primarily in the intracondylar notch, which is in proximity to the remnants of the ruptured ACL. Empty liposome-treated animals showed a greater number of IL-17A+ cells than IDO-1-treated animals, and these cells were located primarily within synovium. These results may reflect the systemic shift in T lymphocyte profile, which translates to a shift in local immune cell phenotype tissue infiltration. Follow-on studies with double-staining to confirm T cell lineage of these CTAL4^+^ and IL-17A^+^ cells are needed.

Articular cartilage loss is a feature of mid- and late-stage PTOA.^7^ Preceding loss, pathologic changes to cartilage structure are detectable with advanced imaging. In this study, contrast-enhanced μCT imaging was used to quantify the thickness and areal surface roughness of distal femoral articular cartilage as a function of injury and treatment. In addition to sex-based differences in cartilage thickness, treatment effects were observed for both male and female rats. Female rats treated with IDO-1-loaded liposomes showed increased AC.Th of the lateral femoral condyle and trochlea compared to female rats treated with empty liposomes, and increased AC.Th of the whole distal femur compared to both female rats treated with empty liposomes, and their own (uninjured) contralateral controls. Male rats treated with IDO-1-loaded liposomes showed increased medial femoral condyle AC.Th when compared to their (uninjured) contralateral controls, but do differences compared to male rats treated with empty liposomes. Despite sex-based differences, articular cartilage areal surface roughness (AC.Sa) was similar between IDO-1-treated and empty liposome-treated animals across all compartments. Partain, et al. observed that intra-articular administration of an IDO-1/Galectin-3 fusion molecule resulted in anti-inflammatory and analgesic effects following joint trauma, but no significant effects with respect to histologic scores of arthritis severity.^49^ It is entirely possible that IDO-1 possesses potent anti-inflammatory and anti-nociceptive effects, but lacks the ability to protect joint tissue structures. Significantly more follow-on studies would be required to substantiate this hypothesis, however.

This study is not without its limitations. First, this study used a rodent model of ACL rupture-induced PTOA. Preclinical models, while integral to discovery-based research, require replication and translation to animals higher in phylogeny to determine the significance of findings to human health. Standardized structural histology, such as that required by OARSI scoring, was not obtained to assess the effect of IDO-1 treatment on PTOA severity. Instead, we used CE-μCT-based measurements of articular cartilage thickness and surface roughness. While these measures provide important three-dimensional structural information, implementation of OARSI scoring could have provided an additional dataset. Specifically, histologic analyses could have provided context for the CE-μCT-based findings of increased AC.Th associated with IDO-1-loaded liposome treatment. Without histology, it is not possible to determine whether the increased AC.Th was a degenerative, hypertrophic response of the cartilage, a mitigation of cartilage thinning, or a transient response with no significant effect on OA outcomes. Accordingly, this experimentation has been started and is ongoing. Further, the CE-μCT-based assessment of PTOA outcomes was limited to a single, early endpoint (2 weeks). While articular cartilage and subchondral bone changes may be detectable at this early endpoint, mid- and late-term follow-up provide important context and trends regarding the severity and progression of PTOA.

## CONCLUSION

Joint trauma leads to dysregulated tryptophan metabolism, which may be a viable target in the treatment and/or prevention of PTOA. Intra-articular administration of IDO-1-loaded liposomes was associated with a reduction in synovial fluid concentrations of tryptophan, IL-1β, and TNF-α. Intra-articular IDO-1 therapy was also associated with a shift in circulating lymphocytes favoring a higher number of T_regs_ compared to T_h_17 cells. Similarly, intra-articular IDO-1 therapy increased the number of CTLA4^+^ cells and reduced the number of IL-17A^+^ cells. IDO-1-treated animals displayed higher mean articular cartilage thickness at a two-week endpoint, but more investigation is needed to determine if this represents a chondroprotective effect versus pathologic remodeling of articular cartilage.

